# Mutational footprint of platinum chemotherapy in a secondary thyroid cancer

**DOI:** 10.1101/2022.03.14.484002

**Authors:** Julia Schiantarelli, Theodora Pappa, Jake Conway, Jett Crowdis, Brendan Reardon, Felix Dietlein, Julian Huang, Darren Stanizzi, Evan Carey, Alice Bosma-Moody, Alma Imamovic, Seunghun Han, Sabrina Camp, Eric Kofman, Erin Shannon, Justine A. Barletta, Meng Xiao He, David Liu, Jihye Park, Jochen H. Lorch, Eliezer M. Van Allen

## Abstract

Although papillary thyroid carcinoma (PTC) is the most frequent endocrine tumor with a generally excellent prognosis, a patient developed a clinically aggressive PTC eleven years after receiving platinum chemotherapy for ovarian endometrioid adenocarcinoma. Germline and somatic analyses of multi-temporal and multi-regional molecular profiles indicated that ovarian and thyroid tumors did not share common genetic alterations. PTC tumors had driver events associated with aggressive PTC behavior, an *RBPMS-NTRK3* fusion and a *TERT* promoter mutation. Spatial and temporal genomic heterogeneity analysis indicated a close link between anatomical locations and molecular patterns of PTC. Mutational signature analyses demonstrated a molecular footprint of platinum exposure, and that aggressive molecular drivers of PTC were linked to prior platinum-associated mutagenesis. This case provides a direct association between platinum chemotherapy exposure and secondary solid tumor evolution, in specific aggressive thyroid carcinoma, and suggests that uniform clinical assessments for secondary PTC after platinum chemotherapy may warrant further evaluation.

## Introduction

Many recent studies have discussed platinum mutagenesis that are observed in tumors being directly treated with these agents as well as in hematologic secondary malignancies^1–6^. However, data supporting a direct clinical association of platinum exposure leading to genetic drivers promoting secondary solid tumorigenesis is currently lacking. A 70-year-old female was diagnosed with ovarian endometrioid adenocarcinoma in 1999, treated with oophorectomy, hysterectomy and chemotherapy (intravenous carboplatin, taxol and topotecan) in 1999, followed by intraperitoneal and intravenous cisplatin and gemcitabine in 1999-2002. In 2002, she experienced a recurrence managed with radiation therapy (3850 cGy intracavitary brachytherapy). Eight years later (2010), she was diagnosed with papillary thyroid carcinoma (PTC), for which she underwent total thyroidectomy and central lymph node dissection. Histopathologic evaluation revealed classic PTC with extensive vascular invasion, extrathyroidal extension and involvement of 3 perithyroidal lymph nodes. Despite radioactive iodine ablation (150mCi in 2010) and longterm thyroid stimulating hormone (TSH) suppression therapy with levothyroxine, she experienced locoregional recurrence on the left lateral neck twice, 4 and 8 years later, treated surgically on both occasions (Figure 1A). Tissue samples from ovarian tumor (n=1), primary PTC from initial thyroidectomy (n=10), and two recurrent PTC excisions (n=1 each) were submitted for molecular profiling (Methods). The locations of tissue samples from primary PTC thyroidectomy, as documented in pathology reports and with approximate relative positions, are depicted in Figure 2A.

**Figure 1.**
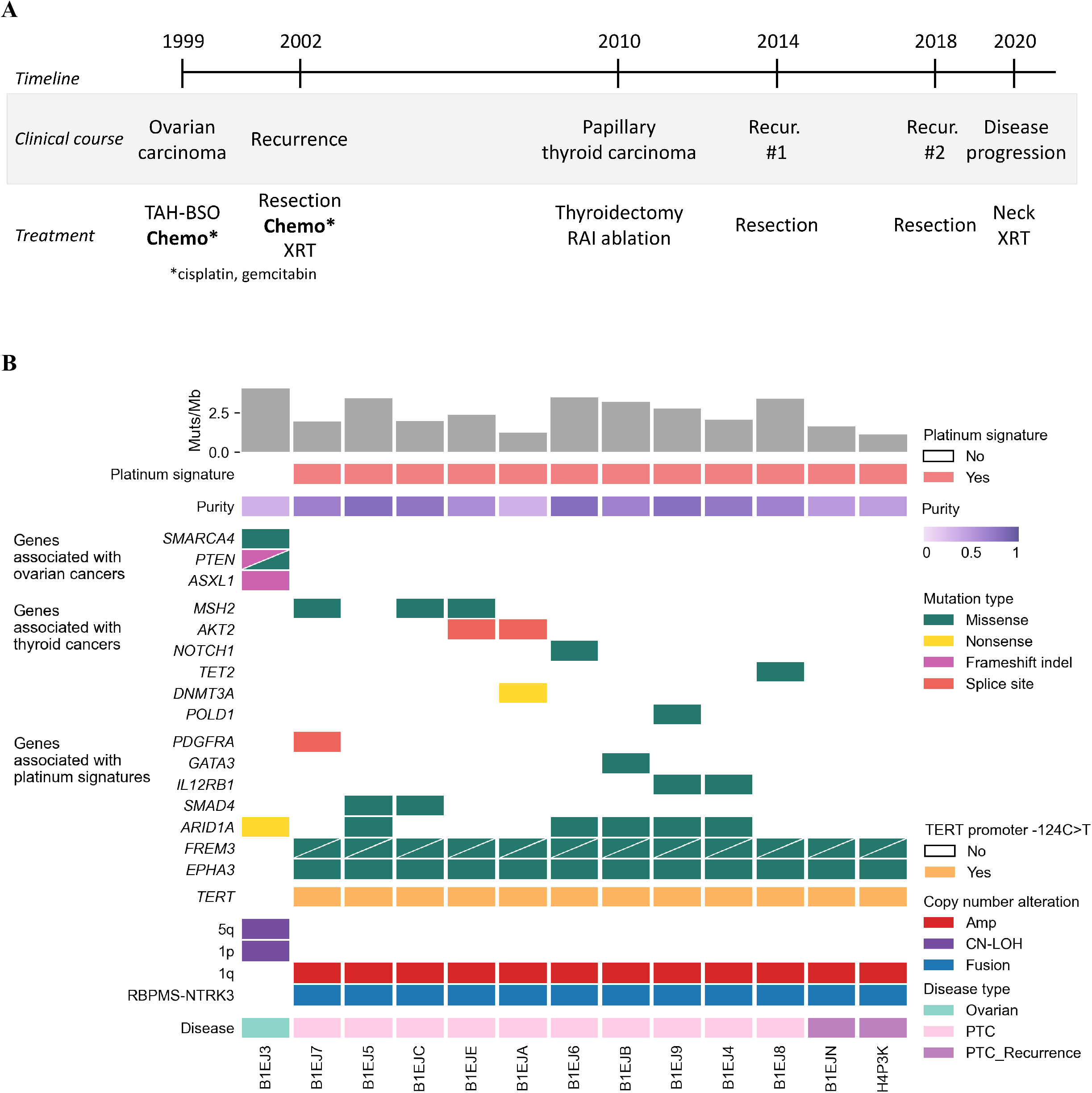
Genomic landscape of the patient’s ovarian and PTC. **A)** Timeline of sample collection, the patient’s clinical course, and treatment history identified in ovarian (1999-2002) and thyroid cancer (2010-2020). **B)** The CoMut^42^ plot illustrates select single nucleotide and insertion/deletion events, as well as CNA, *RBPMS-NTRK3* RNA fusion and *TERT* promoter status. The plot also includes the mutation burden, presence of the platinum signature determined by deconstructSigs and SigProfiler, and tumor purity as determined by ABSOLUTE. Each row represents the mutation or copy number status for the indicated gene, and each column represents a unique tumor sample (Ovarian or PTC samples). Two mutations in the same gene are represented by triangles. CN-LOH indicates copy-neutral loss of heterozygosity.

**Figure 2.**
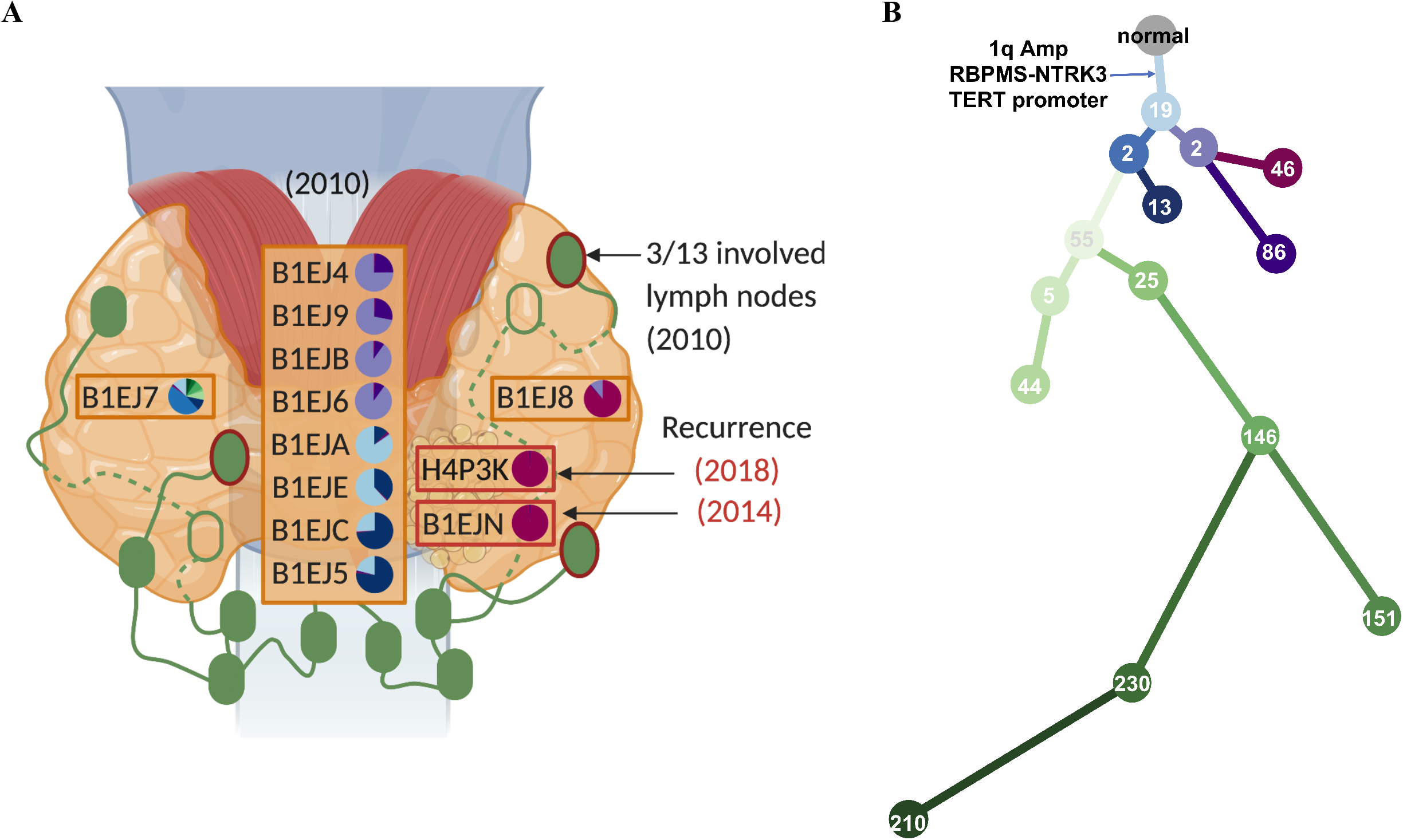
The spatial and temporal tumor heterogeneity in PTC. **A)** A map of the PTC sample locations collected during the initial 2010 thyroidectomy (3 orange boxes with 10 samples) and subsequent surgeries for loco-regional recurrence in 2014 and 2018 (2 red boxes with 2 samples). Three lymph nodes (green ovals with red borders) were positive for the spread of the tumor. Green empty ovals represent 2 lymph nodes posterior to the thyroid. Pie charts represent the fraction of each subclone found in each sample with a common ancestor existing in all PTC samples. **B)** A phylogenetic tree of each subclone represented in the pie charts of Figure 2A. Number in the circle indicates the number of variants assigned to each subclone. The clonal events shared by all PTC samples were annotated in the tree.

## Results

### Molecular origins and evolution of primary ovarian and secondary papillary thyroid cancers

Prior intratumoral heterogeneity (ITH) studies have revealed considerable variations in genetic makeup in tumors across anatomic locations and disease stages^7^, which we hypothesized may inform molecular origins and evolution of this secondary PTC given the aggressive course and prior clinical context. We evaluated multi-regional and multi-temporal samples (total of 12 thyroid samples and 1 ovarian sample) to interrogate genetic makeup of ovarian, primary and recurrent PTC samples. Along with an ovarian cancer sample collected in 1999, 10 samples were collected from different locations in 2010 thyroidectomy and 2 samples from surgeries for PTC recurrence in 2014 and 2018 (Figure 1A). Germline analysis did not identify any known pathogenic genetic alteration associated with ovarian or thyroid cancer or cancer-related genetic syndrome. Comparison of somatic genomic features, including mutations and copy number alterations (CNAs) of the ovarian cancer and PTC samples, indicated that ovarian and thyroid tumor did not share common genetic alterations and originated from genetically distinct tumorigenic events (Figure 1B and S1). The ovarian cancer harbored canonical somatic mutations (e.g. *PTEN, SMARCA4* mutations), whereas PTC tumor harbored driver events including *RBPMS-NTRK3* fusion and a *TERT* promoter mutation, both associated with aggressive PTC behavior (Figure 1B and Table S2)^8,9^.

To explore the evolutionary relationship between tumor foci in these multi-regional and multi-temporal thyroid samples, we clustered mutations to subclones and built a phylogenetic tree representing the relationships between those subclones (Figure 2A-B: Methods). All subclones in PTC samples shared an 1q amplification and canonical driver events implicated in PTC oncogenesis (*RBPMS-NTRK3* fusion and a *TERT* promoter mutation) (Figure 2B and Table S3). We observed four distinct phylogenetic groups across all PTC samples with varying degrees of subclones. Eight samples fell into one of two phylogenetic branches based on their clonal architecture: B1EJA, B1EJE, B1EJC, and B1EJ5 were dominated by the most recent common ancestor of all PTC subclones and a closely related descendent (Figure 2A-B: blue branches); B1EJ4, B1EJ9, B1EJB, and B1EJ6 samples were dominated by a different descendent of the most recent common ancestor (Figure 2A-B: purple branches). Anatomically, these samples were spatially near each other within same phylogenetically defined groups (Figure 2A-B: matching color in pie chart and tree). One site (B1EJ7) had the most diverse subclones (Figure 2A-B: green branches). The two samples that corresponded to loco-regional recurrences in 2014 and 2018 (B1EJN and H4P3K, respectively) shared similar dominant subclones with the PTC sample (B1EJ8) from 2010 thyroidectomy. Sample B1EJ8 was from an area of tumor that was present at the thyroidectomy resection margin and was in closest spatial proximity to recurrences (Figure 2A-B: red branches), supporting the relationship between pathologic findings and molecular spatial patterns of recurrent thyroid carcinoma.

### Cisplatin mutational signature was present in primary and recurrent PTC two decades after chemotherapy

To examine sources of mutagenesis underlying mutation patterns observed in this PTC, we performed mutational signature analysis on the ovarian cancer and PTC samples (Methods). Both ovarian and PTC tumors harbored the ubiquitous clock-like (SBS1 or SBS5) mutational signature (Figure S2–S6 and Table S4). Only the PTC samples, obtained 11-19 years after exposure to platinum chemotherapy, had evidence of platinum mutational signatures (SBS31 or SBS35; Figure S2–S6 and Table S4).

We then calculated the likelihood of observing a mutation in the specific trinucleotide context induced by a specific signature (Methods and Figure 3, S5-S6). An example of a missense mutation mostly attributed to the clock-like (SBS1 or SBS5) signature was an A[C>T]G alteration in *NOTCH1* (Figure 3A-B). Additional mutations related to the platinum chemotherapy signature across the PTC samples were identified in genes such as *EPHA3*, *SMAD4*, and *GATA3* (Figures 1B, 3A-B). The clonal c.-124C>T *TERT* promoter mutation is an established driver of aggressive PTC^9^ and was found in a mutational context, C[C>T]T, characteristic of the platinum chemotherapy signature (Methods and Figure 3A-B, S5–S6). This mutation was present in all PTC tumor samples (Figure 1B and S7) and thus links an event driving the thyroid cancer pathogenesis to treatment exposure that occurred decades previously^1,2^.

**Figure 3.**
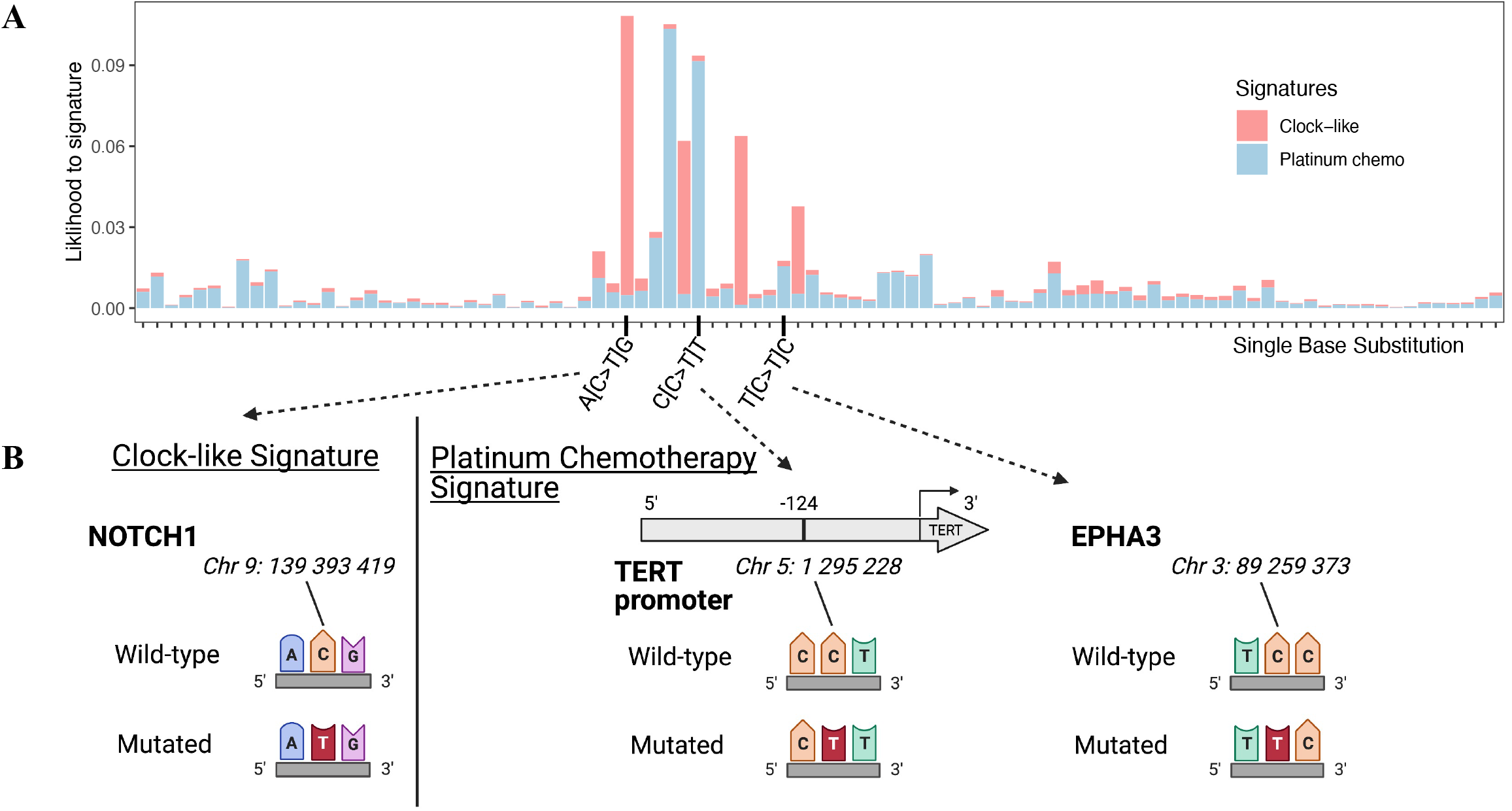
Single base substitution attribution to platinum chemotherapy signature. **A)** The y-axis indicates the likelihood of observing the single base substitution induced by a specific signature utilizing SigProfiler. The x-axis indicates single base substitution in 96 trinucleotide contexts. SBS1 and SBS5 were summarized as Clock-like signatures. SBS31 and SBS35 were indicated as Platinum chemotherapy signatures. **B)** Nucleotide context with single base substitution changes in *NOTCH1, TERT* promoter and *EPHA3* genes.

## Discussion

Several recent studies discussed the chemotherapy mutational signatures that have been detected in metastatic tumors or in secondary hematological malignancies^1–6^. However, molecular origins and evolution of secondary solid tumors in patients previously exposed to platinum-based chemotherapy remain to be elucidated, although there are paradigms of chemotherapy induced mutagenesis leading to drug resistant clones and impacting clinical outcomes^1,2^. Here, molecular profiling of the PTC at different timepoints and locations revealed significant intratumoral heterogeneity and genetic alterations distinct from ovarian cancer of the same patient (Figure 1 and S1). Thyroid cancer is typically considered largely homogeneous at the molecular level, with the TCGA study proposing two major classifications: BRAF-like and RAS-like^10^. Only a small body of literature supports the presence of concomitant mutations, heterogeneous presence of driver mutations (such as *BRAF^V600E^*), and discordant profile of primary and metastatic PTC^11–14^. By exploring evolutionary relationships across multi-regional and multi-temporal samples from the same patient, we illustrated the relationships between different subclones, and branches of samples with distinct subclones that were associated with the anatomical locations of collected samples (Figure 2 and Table S3). As the pathology report noted positive margins on excision of patient’s tumor during 2010 thyroidectomy, it is possible that loco-regional recurrences arose by remnant cancer cells escaping radioactive iodine ablation. No *BRAF* or *RAS* mutation was identified in primary and recurrent PTC samples. Instead, all PTC samples harbored *RBPMS-NTRK3* fusion, *TERT* promoter c.-124C>T mutation and 1q amplification, which suggests that these genetic alterations are closely linked with this PTC’s pathogenesis and aggressive features (Figure 1 and 2). NTRK altered PTC is rare, comprising under 2% of cases, and is characterized by multinodular growth, prominent fibrosis, extensive lymphovascular spread and high risk of recurrence and metastatic disease^8^, which is consistent with this patient’s tumor pathology and behavior. Similarly, *TERT* promoter mutation, enriched in less well differentiated and anaplastic thyroid carcinomas^9^, and 1q amplification are also associated with higher disease stage^10^.

Platinum chemotherapy mutational signatures were observed in all PTC samples, a footprint present 19 years after chemotherapy exposure (Figure S2–S6 and Table S4). Tumor location and primary *vs* recurrent site did not impact the degree of platinum mutational signatures observed; however, we demonstrated that c.-124C>T *TERT* promoter single base substitution in C[C>T]T context was mostly attributed to platinum mutational signature (Figure 3 and S5–S6). While ionizing radiation is a well-established risk factor for PTC, and gene fusions (in particular *RET-PTC* rearrangements) and copy number alterations are enriched in radiation induced PTC^15,16^, no such link has been recognized for chemotherapy. Yet, in a study of 12,547 childhood cancer survivors, treatment with alkylating agents was associated with increased PTC risk, beyond the relative risk that could be attributed to prior ionizing radiation therapy^17^. This PTC case harbored uncommon genetic patterns with *NTRK* fusion and 1q amplification. Taken together and with prior knowledge of how chemotherapy may induce DNA damage and breakpoints^18,19^, these findings provide a mechanistic hypothesis for how platinum mutagenesis might have induced a *TERT* promoter mutation and contributed to aggressive histopathological and clinical course of this patient’s PTC.

This study may offer a mechanistic explanation for elevated thyroid cancer risk in patients after platinum chemotherapy exposure^2,17^, who may benefit from increased awareness and lower threshold to screen for secondary PTC. For those exposed to platinum chemotherapy who do later develop thyroid cancers, looking for rare but prognostically significant driver events may be informative. The American Thyroid Association guidelines^20^ do not specifically address thyroid cancer screening in chemotherapy treated patients, and there is limited data on the impact of chemotherapy on thyroid tumorigenesis. Future studies of a larger cohort of thyroid cancer patients with exposure to chemotherapy for a previous cancer will be necessary to determine whether there is a larger pattern of platinum chemotherapy induced driver mutations explaining increased incidence of thyroid cancer seen in this population.

## Methods

### Sample preparation and sequencing

Written informed consent was obtained by the patient for participation in the study under the Dana-Farber Cancer Institute’s Institutional Review Board 09-472. DNA extraction, library preparation, and WES/WGS were performed for samples as previously described^21^. Slides were cut from FFPE tissue blocks and microdissected for tumor-enriched tissue. DNA and RNA extraction were performed using QIAGEN AllPrep FFPE DNA/RNA extraction kit. Germline DNA was obtained from the peripheral blood sample. Libraries were constructed, hybridization and capture were performed with Illumina’s Rapid Capture Exome Kit for WES, and then sequenced with Illumina HiSeq as previously described^22^. cDNA library synthesis and capture were performed using the Illumina TruSeq RNA Access Library Prep Kit. Flowcell cluster amplification and sequencing were performed with Illumina sequencers. Each run was a 76bp paired-end and Broad Picard Pipeline was used for de-multiplexing and data aggregation.

### Germline analysis

Germline WES data was analyzed using DeepVariant^23,24^ to search for alterations with clinical significance (pathogenic or likely pathogenic) and known variants associated with germline cancer predisposition. DeepVariant is a deep convolutional neural network, based on the inception framework, trained to identify inherited variants from read pileup pseudo-images. We ran DeepVariant using recommended settings (https://github.com/google/deepvariant).

### WES analysis

Variants were called from the WES data with the customized version of the CGA pipeline (https://github.com/broadinstitute/CGA_Production_Analysis_Pipeline) (Table S1-S2). Quality control was performed by estimating contamination with ContEST^25^ and utilizing Picard Multiple Sequence Metrics. Copy number alterations (Figure S1) were called with GATK CNV^26^ and Allelic CapSeg^27^, while Mutect1^28^ and Strelka^29^ were used to call SNVs and indels. Mutect2 was used to confirm Strelka indel calls. Somatic mutation calls were filtered for FFPE and 8-OxoG sequencing artifacts using GATK FilterByOrientationBias^30^ and further filtered against a panel of normals of similarly prepared samples. Finally, we utilized ABSOLUTE^31^ to determine allelic copy number, tumor purity and tumor ploidy. A union set of mutations from all samples was generated and force calling was performed to assess the variant allele fraction of each mutation within each sample. The cancer cell fraction (CCF) of mutations were calculated via ABSOLUTE^31^. To reconstruct the clonal architecture of each thyroid tumor, we used the PhylogicNDT^32^ cluster module to determine the number of subclones and assigned mutations to each subclone (Table S3). Then, we built the phylogenetic trees based on PhylogicNDT subclones, using the BuildTree module. Multiple trees can be constructed for any given set of subclones, and we used the maximum likelihood tree in our analyses.

### Mutational signature analysis

Mutational signatures were determined using deconstructSigs^33^ as well as SigProfiler^34^ with COSMIC v3 signatures^34^ as the reference. A signature cutoff was set 0.08 for deconstructSigs and exome parameter was True and SBS96 context was used for SigProfiler (Table S4 and Figure S2–S6). Both methods identified clock-like signature (SBS1 or SBS5) and platinum chemotherapy signature (SBS31 or SBS35) as the two dominant signatures contributing to our tumor samples. Specifically, both methods identified platinum chemotherapy signature in all thyroid cancer samples but not in the ovarian cancer sample (Figure 1B and S2–S4). We calculated the likelihood of observing a mutation in the specific trinucleotide context induced by a specific signature^2^, and identified the mutation candidates that are more likely to be induced by either clock-like signature or platinum chemotherapy signature utilizing decomposed SigProfiler results (Figure 3A-B and Figure S3–S5). Since the clock-like signature and platinum chemotherapy signature are two dominant signatures decomposed from SigProfiler results, we also calculated the relative contribution of either signature (clock-like vs platinum chemo) given the specific trinucleotide context (Figure S6). The presence of a TERT promoter mutation (c.-124C>T), known to be associated with more aggressive PTC^35^, was reported on review of the patient’s medical records documenting the results of an independent genetic platform. While we mapped mutations induced by a specific mutational signature, we also searched for the presence of *TERT* promoter mutations (i.e. c. −124C > T (C228T) and c. −146C > T (C250T)). Manual inspection with Integrative Genomics Viewer (IGV)^36^ was performed on associated WGS samples from this patient and confirmed the presence of the c.-124C>T *TERT* promoter mutation in all PTC samples (primary and recurrent) but not in the ovarian cancer sample (Figure S7). As promoter regions do not overlap with WES target regions, we used available low depth WGS instead of WES data to confidently detect the *TERT* promoter mutation.

Formalin fixation and storage is known to cause DNA fragmentation and cytosine deamination, and studies have shown “C:G > T:A” artifacts with low allelic frequency (<5%)^37,38^. Platinum chemotherapy signatures also share “C:G > T:A” substitutions in specific trinucleotide context, so we also performed mutational signature analysis with high allelic frequency (5%) and low allelic frequency (<5%) mutations separately even after FFPE filtering steps that were already performed in our WES analysis. We still detected platinum mutational signatures in all PTC samples with high allelic frequency mutations, whereas in low allelic frequency mutation analysis the platinum mutational signatures were not as strongly evident. (Figure S2–S6 and Table S4).

### RNAseq analysis

We utilized STAR and RSEM to quantify gene and isoform abundances from RNA-seq data. Probabilistically weighted alignments and gene abundance estimates were generated, and the methods output an expected read count distribution. All RNA samples were sequenced together in the same batch, and we set benchmarks for the maximum intergenic, intronic, and rRNA rates. Based on this, we excluded samples that did not meet these criteria. We utilized three fusion callers: STAR Fusion^39^, ChimPipe^40^, and FusionCatcher^41^. These callers capture split-reads and discordant paired-end reads, map them to reference, filter fusion predictions, and annotate the calls. We looked for fusions identified in each sample by two or more of the three callers (Figure 1B).

## Acknowledgments

This work was supported by NCI F31CA239347 (to J. Conway.), NIH T32 GM008313 and NSF GRFP DGE1144152 (to M.X.H.), NIH T32 5T32HL007609–33 (to T.P.), NIH R37 CA222574 (to J.P. and E.M.V.), NIH R01 CA227388 (to J.P. and E.M.V.), and Damon Runyon Clinical Investigator Award (E.M.V.). We are incredibly grateful for the patient who made this study possible.

## Author Contributions

J.S. and T.P. contributed equally to this work. E.M.V., J.H.L., and J.P. jointly supervised this work. J.S., T.P., J.P., J.Conway, J.Crowdis, B.R., F.D., A.B., A.I., S.H., S.C., E.K., M.X.H., D.L., J.H.L., and E.M.V. contributed to genomic analysis and interpretation of results. J.H., D.S., E.C., E.S., J.P., J.H.L., E.M.V. contributed to sample and clinical information collection. J.A.B. contributed to the review of pathological findings. J.S., T.P., J.P., J.H.L., and E.M.V. wrote the manuscript and prepared the figures, which all authors reviewed.

## Data availability

All reasonable requests for raw and analyzed data and materials will be promptly reviewed by the senior authors to determine whether the request is subject to any intellectual property or confidentiality obligations. Patient-related data not included in the paper may be subject to patient confidentiality. Any data and materials that can be shared will be released via a material transfer agreement. All analyzed sequencing data are provided in the Supplementary Information.

---

**Figure S1.**
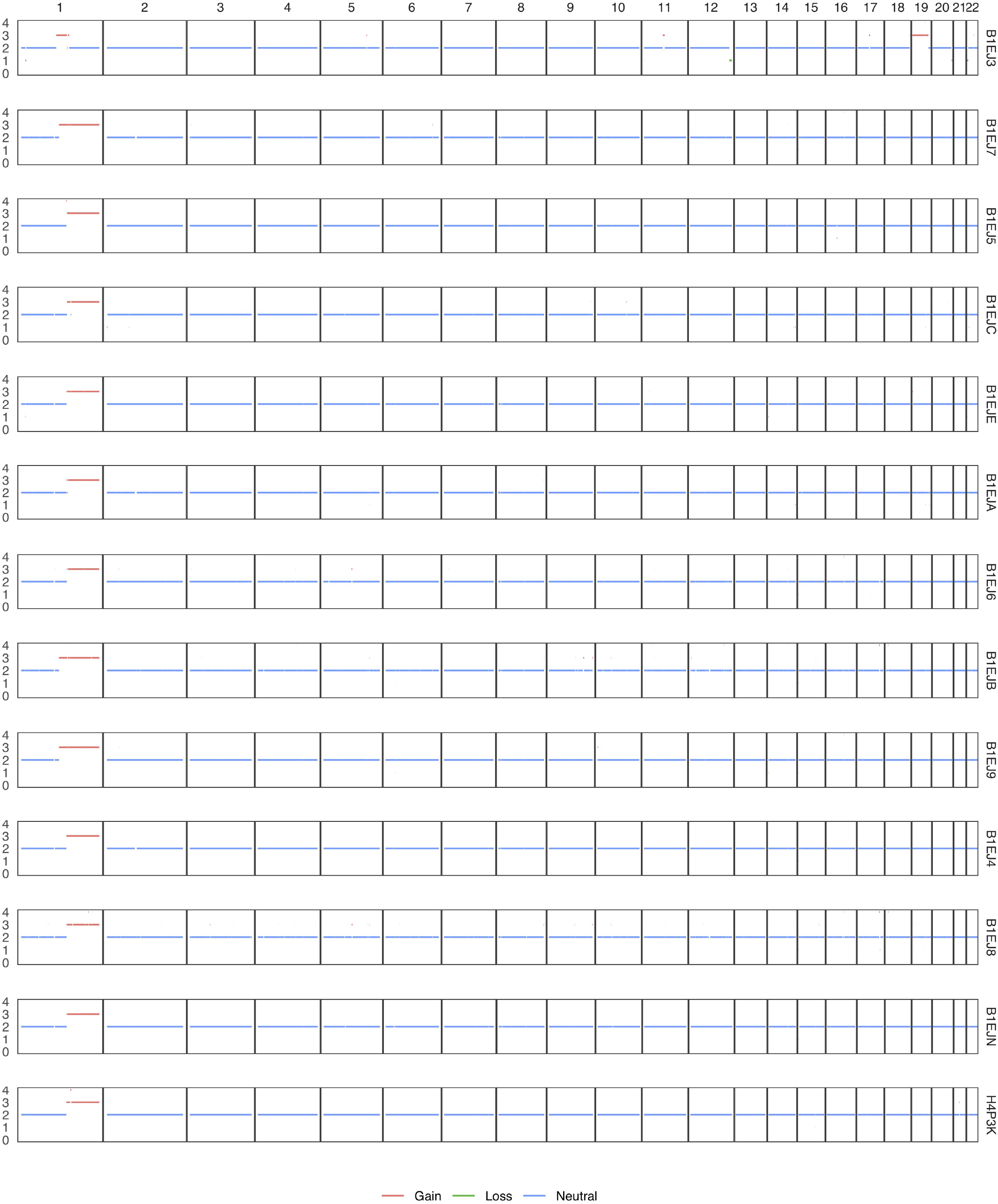
Copy number profile of all samples. The y-axis indicates the total copy number of Ovarian (B1EJ3) and all PTC samples. The x-axis indicates each chromosome.

**Figure S2.**
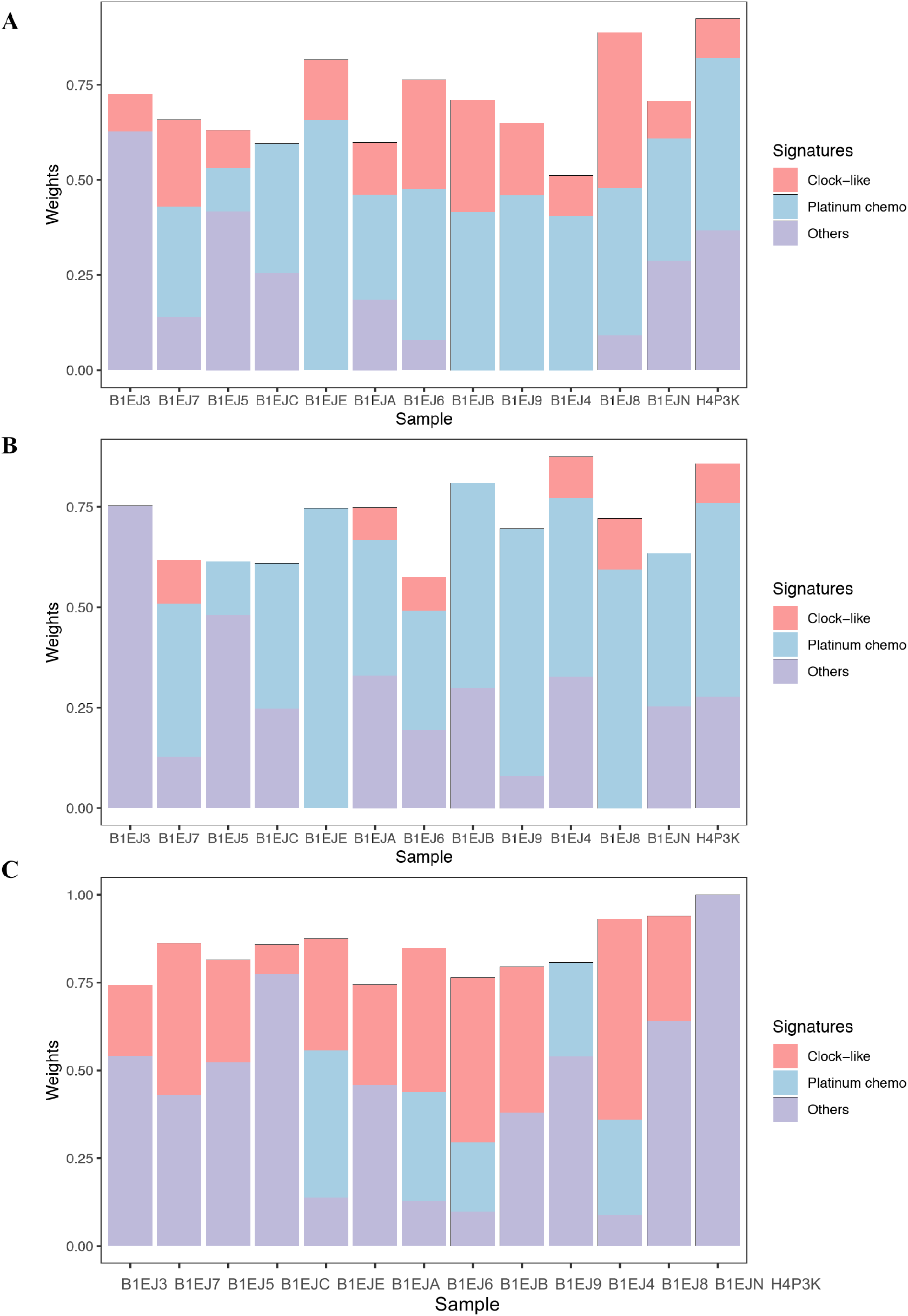
Mutational signatures detected by deconstructSigs. **A)** The y-axis indicates the weights of signatures contributing to Ovarian (B1EJ3) and all PTC samples utilizing deconstructSigs. SBS1 and SBS5 were summarized as Clock-like signatures. SBS31 and SBS35 were indicated as Platinum chemotherapy signatures. All other signatures were shown as Others. **B)** Signatures detected by all mutations >= 0.05 allelic fraction. **C)** Signatures detected by all mutations < 0.05 allelic fraction.

**Figure S3.**
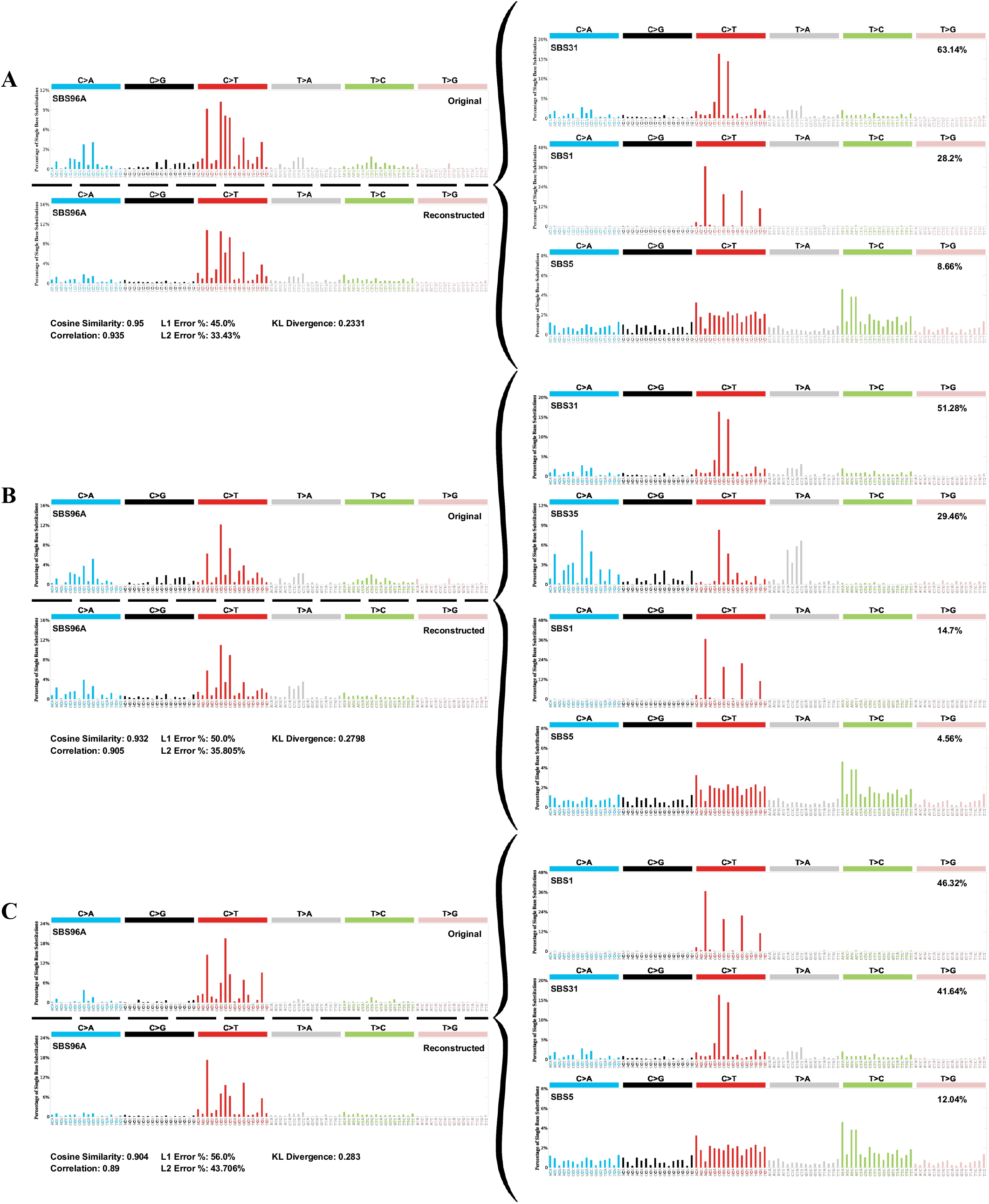
Mutational signatures decomposed by SigProfiler. **A)** Left top panel shows the original signature and left bottom panel shows the reconstructed signature. Right panels show the decomposed signatures reference to COSMIC v3. **B)** Signatures detected by all mutations >= 0.05 allelic fraction. **C)** Signatures detected by all mutations < 0.05 allelic fraction.

**Figure S4.**
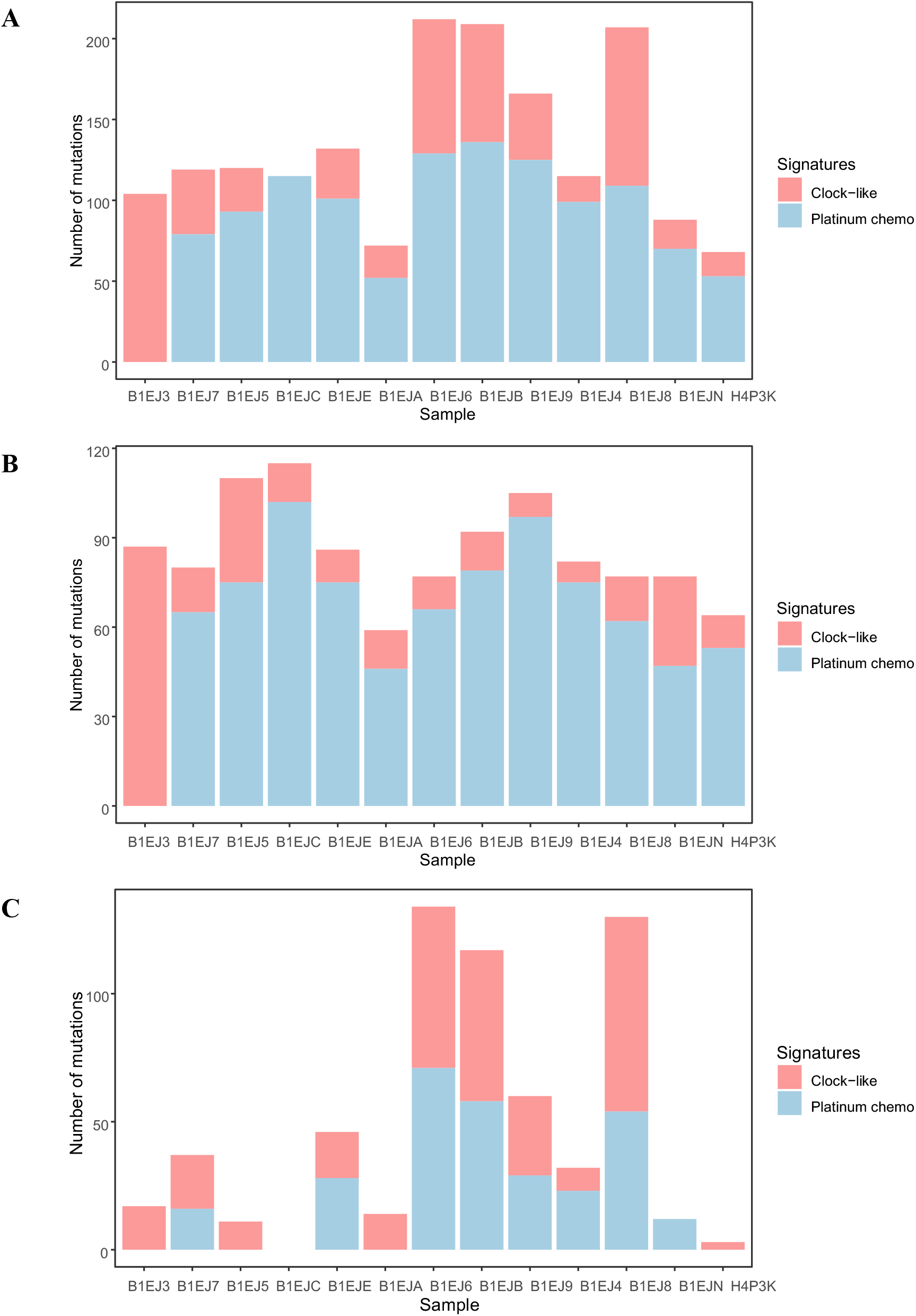
Mutations for each signature detected by SigProfiler. **A)** The y-axis indicates the number of mutations contributed to each signature in Ovarian (B1EJ3) and all PTC samples utilizing SigProfiler. SBS1 and SBS5 were summarized as Clock-like signatures. SBS31 and SBS35 were indicated as Platinum chemotherapy signatures. **B)** Signatures detected by all mutations >= 0.05 allelic fraction. **C)** Signatures detected by all mutations < 0.05 allelic fraction.

**Figure S5.**
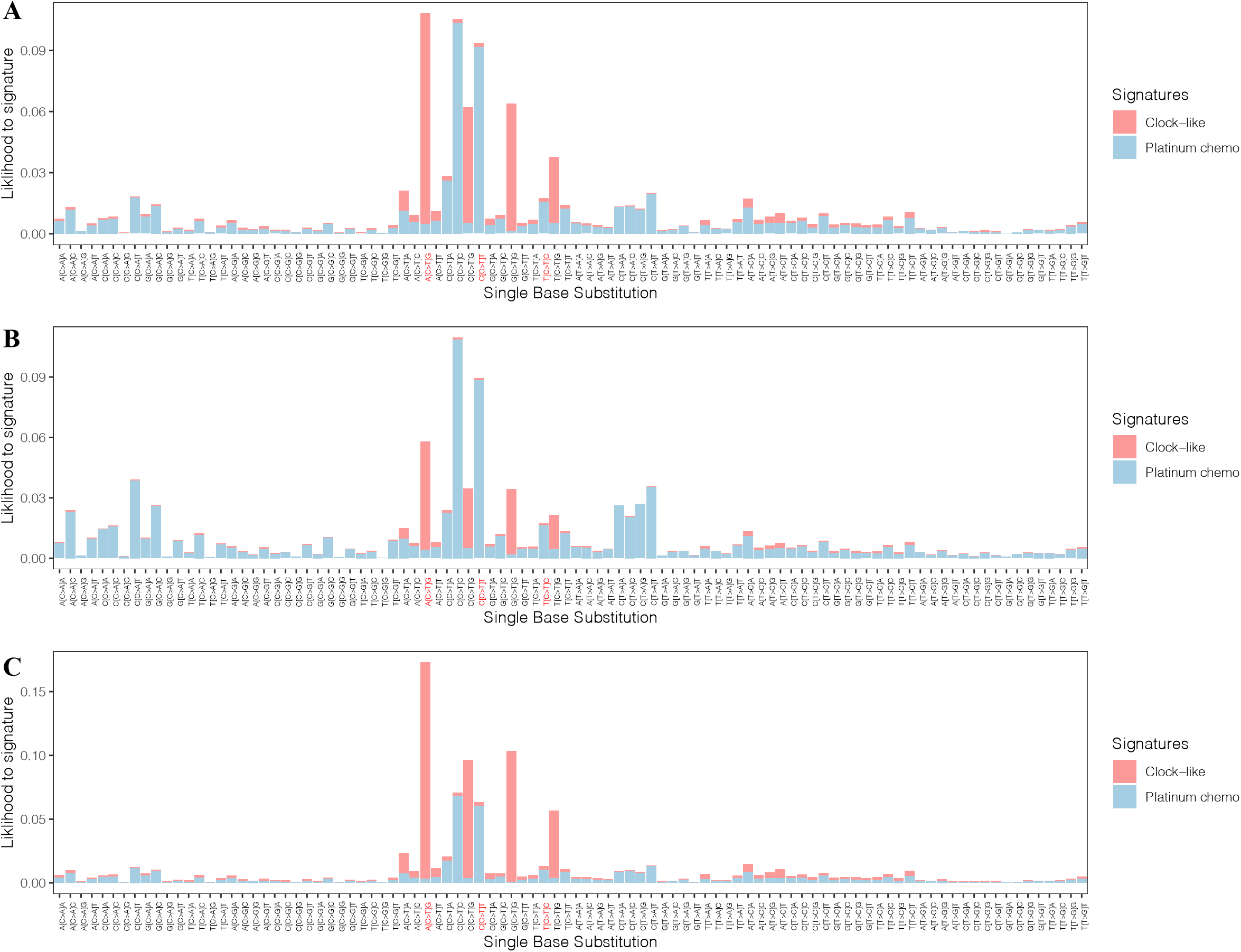
Single base substitution attribution to signatures detected by SigProfiler. **A)** The y-axis indicates the likelihood of observing the single base substitution induced by a specific signature utilizing SigProfiler. The x-axis indicates single base substitution in 96 trinucleotide contexts. Three trinucleotide contexts were highlighted with red based on Figure 3 examples (A[C>T]G for *NOTCH1*, C[C>T]T for *TERT* promoter, and T[C>T]C for *EPHA3*). SBS1 and SBS5 were summarized as Clock-like signatures. SBS31 and SBS35 were indicated as Platinum chemotherapy signatures. **B)** Signatures detected by all mutations >= 0.05 allelic fraction. **C)** Signatures detected by all mutations < 0.05 allelic fraction.

**Figure S6.**
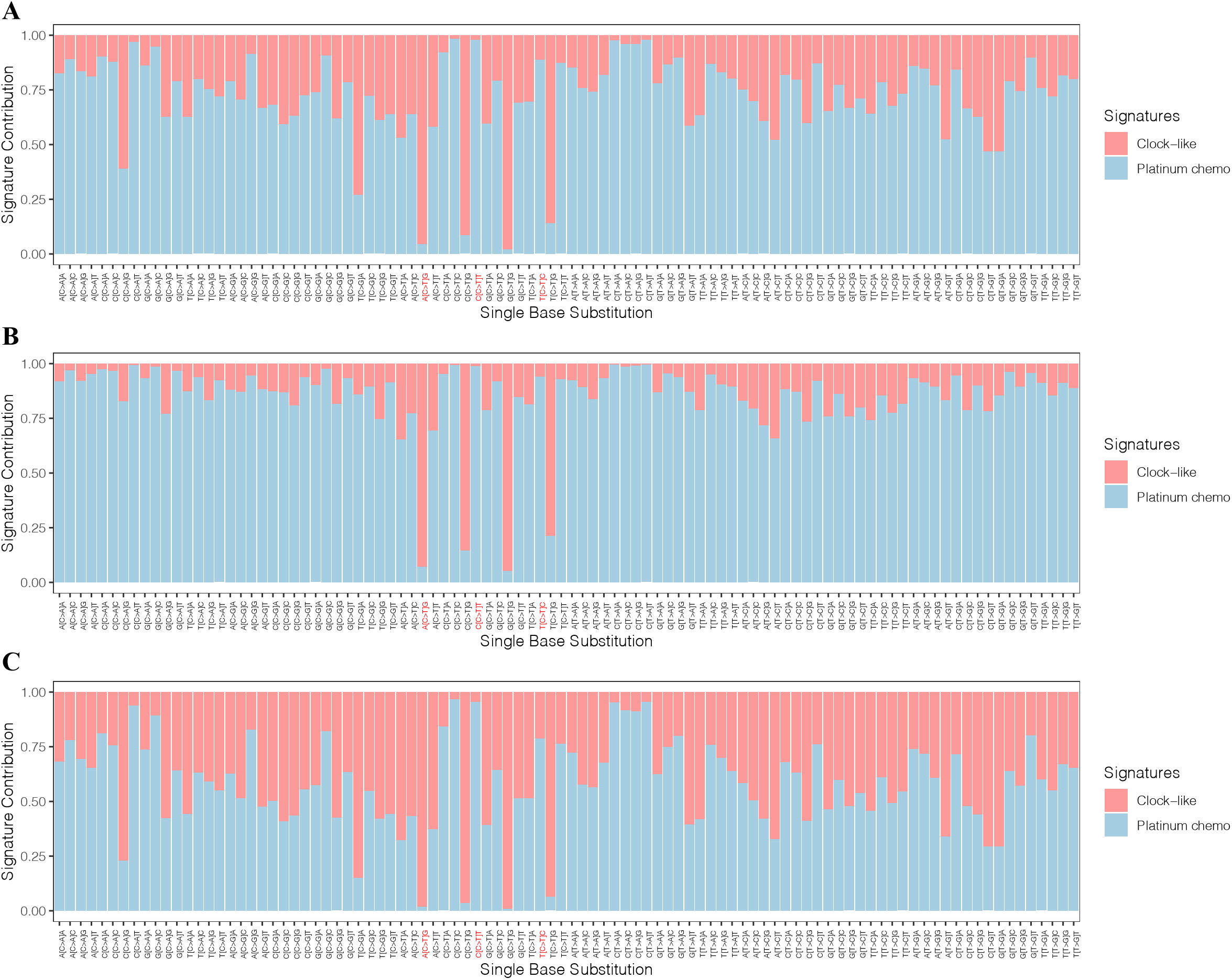
Relative signature contribution by SigProfiler. **A)** The y-axis indicates the relative contribution of either signature (clock-like vs platinum chemo) given the single base substitution utilizing SigProfiler. The x-axis indicates single base substitution in 96 trinucleotide contexts. Three trinucleotide contexts were highlighted with red based on Figure 3 examples (A[C>T]G for *NOTCH1*, C[C>T]T for *TERT* promoter, and T[C>T]C for *EPHA3*). SBS1 and SBS5 were summarized as Clock-like signatures. SBS31 and SBS35 were indicated as Platinum chemotherapy signatures. **B)** Signatures detected by all mutations >= 0.05 allelic fraction. **C)** Signatures detected by all mutations < 0.05 allelic fraction.

**Figure S7.**
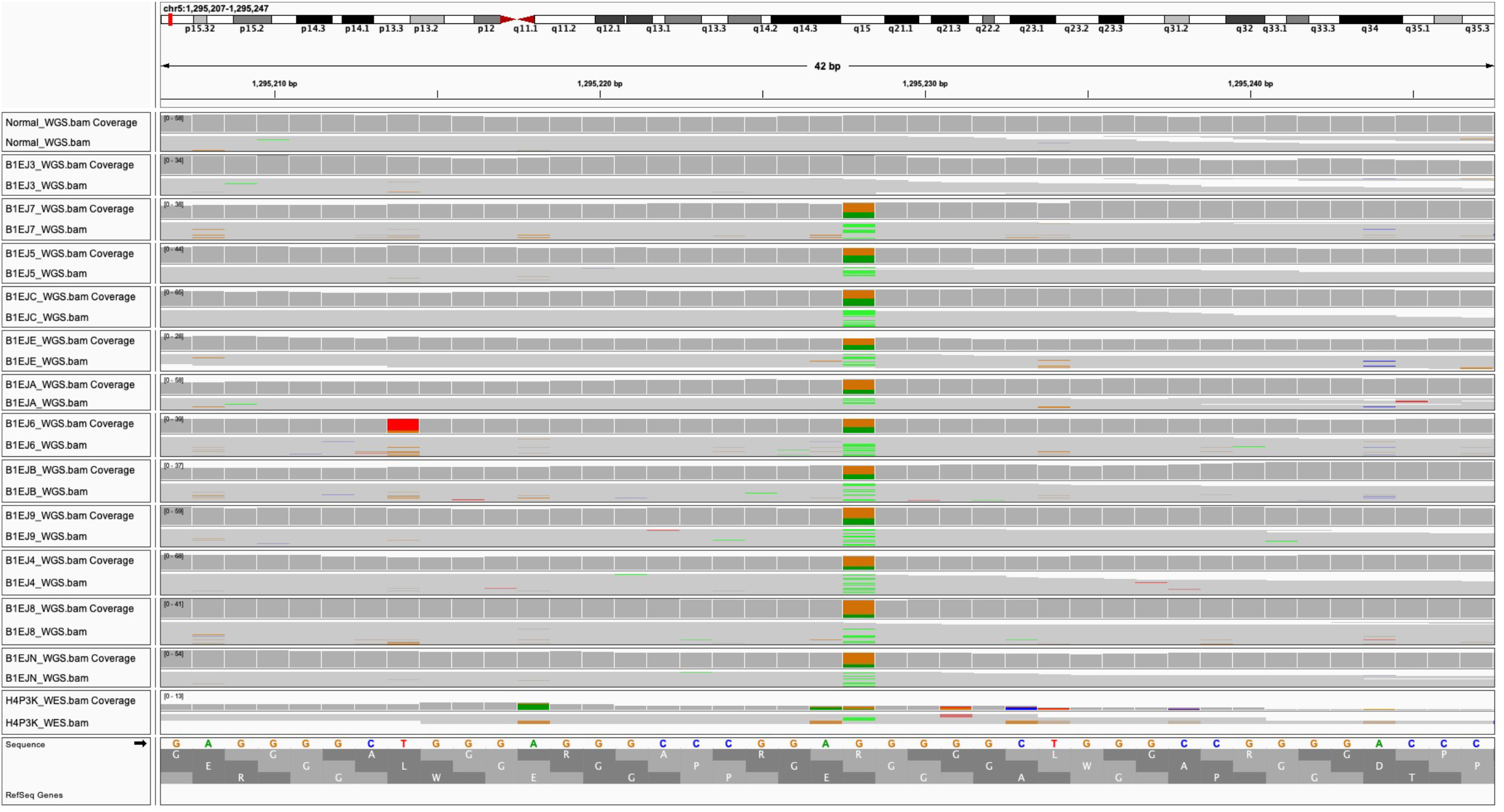
Integrative Genomics Viewer (IGV) snapshot of c.-124C>T *TERT* promoter mutation. Reads are summarized in a coverage plot for each WGS data from germline (Normal), ovarian (B1EJ3) and all PTC samples (except H4P3K which only had WES data). Positions with a significant number of mismatches with respect to the reference sequence at the bottom are highlighted with color bars indicative of both the presence of mismatches (G > A) and the allele frequency (A: green, G: brown). Mutations are only observed in PTC samples not in germline or ovarian samples (top two panels). The reads have been sorted and colored by base.

**Table S1.** Genomic characteristics of all exomes included in study

Uploaded as a separate file

**Table S2.** Union set of mutations in study

Uploaded as a separate file

**Table S3.** PhylogicNDT mutational clustering output

Uploaded as a separate file

**Table S4.** Mutational process as assessed by deConstructSigs and SigProfiler

Uploaded as a separate file

